# RGB color indices as proxy for symbiont cell density and chlorophyll content during coral bleaching

**DOI:** 10.1101/2024.12.20.629333

**Authors:** Erik Francesco Ferrara, Lavinia Bauer, Giulia Puntin, Friederike Bautz, Sibel Celayir, Marie-Sa Do, Frederik Eck, Melissa Heider, Pia Wissel, Angelina Arnold, Thomas Wilke, Jessica Reichert, Maren Ziegler

## Abstract

Coral bleaching, the breakdown of the symbiosis between the coral host and endosymbiotic microalgae, is the main cause of widespread coral reef degradation. Current methods for assessing coral health based on visual appearance, such as the use of color reference cards, are limited by subjective human color perception and low resolution. Digital photography with RGB (Red, Green, Blue) color channel analyses offers a fast, non-invasive, and standardized alternative to estimate physiological parameters. However, the link between coral color and physiological parameters during bleaching may vary depending on the type of stressor. While such approaches are extensively used in plant studies, their application in estimating Symbiodiniaceae cell density and chlorophyll content in corals requires further attention. In this study, we analyzed the correlation between Symbiodiniaceae cell density and chlorophyll content across three coral species (*Acropora muricata*, *Pocillopora verrucosa,* and *Stylophora pistillata*) with 19 color indices derived from the RGB channels currently established as predictors of chlorophyll content in plants. Corals were exposed to three bleaching conditions (acute short-term and chronic long-term heat stress and menthol bleaching) to identify the best color indices for assessing coral health through image analysis. We found that the Red index had the strongest linear correlation with symbiont cell density and chlorophyll content across species (R^2^ up to 0.97), so that relative changes in this color index can be directly interpreted as corresponding changes in tissue parameters. To train a model that predicts symbiont densities of a distinct sample set using the Red index, we found 10 to 12 samples to be sufficient to achieve an accuracy of > 95 % of the models trained on the full datasets. This research contributes to improved image analysis as a reliable and non-invasive tool for monitoring, by providing guidelines for a systematic use of RGB data to interpret coral health.

## Introduction

Coral reefs are among the most biologically diverse and economically valuable ecosystems on Earth, supporting more than 25 % of marine biodiversity and providing valuable ecosystem goods and services, such as protecting coastlines from erosion and storms and representing a natural nursery for many marine species (Ferrario et al., 2014; Moberg & Folke, 1999). Climate change and local stressors threaten the stability of coral reef ecosystems, leading to their global decline (Hughes et al., 2018). Exposure to elevated seawater temperatures above local thermal thresholds disrupts the symbiosis between corals and the endosymbiotic dinoflagellate Symbiodiniaceae, resulting in the loss of the symbiont cells and photosynthetic pigments—a phenomenon known as coral bleaching (Helgoe et al., 2024). During the last decades, the increased frequency and severity of mass coral bleaching events have affected over 70 % of reefs, leading to high global coral mortality (Eakin et al., 2019). In April 2024, the National Oceanic and Atmospheric Administration (NOAA) confirmed the fourth coral bleaching event, highlighting the urgent need to develop effective tools to guide reef management efforts.

To respond to these challenges, a rapid, objective, and non-invasive method to assess coral health and early signs of stress is vital. Visual inspection using waterproof color reference cards first developed by Siebeck et al. (2006) is one of the most commonly used methods to assess the bleaching state in both field surveys and laboratory experiments (Anthony et al., 2008; Conti-Jerpe et al., 2020; Doering et al., 2021; Oladi et al., 2017; Quigley et al., 2020; Rosado et al., 2019; Schoepf et al., 2019). However, color reference cards have been tailored to specific coral species or regions, limiting their broader applicability (Bahr et al., 2020; Bollati et al., 2020; Herrera et al., 2023; Siebeck et al., 2006). Additionally, such methods have limitations related to subjective human color perception, which reduces reproducibility (Parkinson et al., 2016; Siebeck et al., 2006), the capacity to detect small changes in coloration due to the number of color ranks commonly used (Winters et al., 2009), and color gradients within coral fragments (Coles & Brown, 2023).

With the advancement in digital photography, the analysis of high-resolution images has become a promising tool for assessing coral health (Teague et al., 2022). This technique builds on well-established methods used for monitoring the health and growth of terrestrial plants, where red, green, and blue (RGB) color channels of digital images have been successfully used to estimate chlorophyll content, by correlating changes in RGB values with changes in leaf pigmentation (Wang et al., 2014; Widjaja Putra & Soni, 2018; Zhang et al., 2022). Other RGB-derived indices have been tested to improve these correlations and assess plant physiological parameters, such as leaf nitrogen or carotenoid concentration (Kior et al., 2024). In this frame, the RGB channels, centered at approximately 660 nm, 520 nm, and 450 nm wavelengths, respectively, score the coral color on a scale of 0 to 255, with higher values representing lighter color shades. Therefore, variations in channel intensity can be interpreted as changes in specific physiological parameters. For example, Winters et al. (2009) found that changes in red channel values correlated with the chlorophyll content in *Stylophora pistillata*. This suggests that photographic measurements can be used to objectively evaluate the severity of bleaching events based on color channel values.

While color-based assessments of symbiotic status are commonly used (Chow et al., 2016; Ferrara et al., 2024; Reichert et al., 2021; Voolstra et al., 2020), a systematic assessment is needed to determine which RGB indices best capture physiological changes in corals during bleaching and whether these relationships remain consistent across species. Additionally, the correlation between visual appearance and physiological parameters may vary depending on the specific bleaching conditions (Fitt et al., 2000). For instance, different bleaching triggers—such as acute vs. chronic heat stress or chemical agents like menthol—elicit distinct physiological responses, that are likely to affect how RGB values correlate with key metrics such as Symbiodiniaceae cell density and chlorophyll content. Additionally, unlike terrestrial plants, corals present unique challenges for photographic analysis with two main factors potentially interfering with image data interpretation: First, the presence of endolithic algae and pigmentation within the skeleton (Galindo-Martínez et al., 2022) and second, the production of host-derived non-photosynthetic pigments that help reduce light intensity in the tissues in the absence of Symbiodiniaceae cells, a phenomenon referred to as colorful bleaching (Bollati et al., 2020). Therefore, it is essential to refine image analysis techniques specifically for corals to ensure reliable assessments of their health status.

Here, we systematically tested 19 color indices, derived from RGB channels, as proxies for Symbiodiniaceae cell density and total chlorophyll content in *Acropora muricata* (Linnaeus, 1758), *Pocillopora verrucosa* (Ellis & Solander, 1786), and *Stylophora pistillata* (Esper, 1792) exposed to different bleaching conditions. Specifically, we aimed to 1) analyze the correlation between Symbiodiniaceae cell density (and total chlorophyll content) and the color indices, already established in plant studies as good proxies for chlorophyll content, and 2) identify the best predictor of these physiological parameters across different coral species and bleaching conditions, including short-term heat exposure, long-term heat exposure, and menthol bleaching. This approach will be useful for studies aimed at developing strategies to mitigate the impact of global warming on corals by providing a rapid and objective method to assess coral health status. Additionally, we calculated the minimum sample size required to train a high-performance linear model for predicting coral physiological parameters and provided an easy-to-use R script to extract RBG values from coral images that can be used by non-specialists.

## Materials and methods

### Experimental design and study species

Data were collected from three experiments conducted at the *Ocean2100* coral aquarium facility at Justus Liebig University Giessen, Germany, to investigate the correlation between RGB-derived color indices and physiological parameters in corals under different bleaching conditions. Each experiment examined the color-to-physiology correspondence of three common reef-building coral species: *Acropora muricata*, *Pocillopora verrucosa*, and *Stylophora pistillata*.

Prior to the bleaching experiments, all coral fragments were maintained under constant water parameters in tanks connected to an artificial seawater recirculation system. Water temperature was maintained at 26 ± 0.5°C using 300 W heaters feedback-controlled by aquarium computers (GHL Temp Sensor digital, ProfiLux 3 and 4, GHL Advanced Technology GmbH, Germany), and coral fragments of ∼3-4 cm in length were maintained under a 10:14 light:dark photoperiod with light intensity (PAR) of ∼250 ± 30 µmol photons m^-2^ s^-1^ (measured by Apogee Lightmeter, model MQ-510), and salinity of 36. Corals were fed copepods (Calanoide Copepoden, Zooschatz, Germany) three days per week.

### Bleaching protocols

Corals were exposed to three bleaching treatments: short-term heat stress, long-term heat stress, and menthol bleaching. The short-term bleaching experiment was conducted in 40-L tanks equipped with a current pump (easyStream pro ES-28, Aqualight GmbH, Bramsche/Lappenstuhl, Germany) under a light intensity of ∼120 μmol photons m^-2^ s^-1^ (white and blue sunaECO LED, AquaRay by Tropical Marine Centre, United Kingdom). A total of 53 fragments from three coral species were randomly distributed in 6 heat and 6 control tanks with 19 fragments from 8 colonies of *A. muricata*, 18 fragments from 7 colonies of *P. verrucosa*, and 16 fragments from 5 colonies of *S. pistillata*. In the heat tanks, the temperature was increased from 26 °C to 35 °C in three hours, then held at 35 °C for three hours, and then decreased back to 26 °C within two hours. In the control tanks, the temperature was maintained at 26°C (additional details in Ferrara et al., 2024). Physiological parameters were measured 18 h after the start of the heat stress following Voolstra et al. (2020).

For the short-term heat stress experiment, we used the same three species with three colonies per species and six fragments per colony (18 fragments per species, 54 fragments in total). The fragments were distributed among three 40-L tanks, and the water temperature was gradually increased from 26.5 °C to 32.5 °C over 31 days. The temperature was increased daily by 0.5 °C until 29 °C in five days, then by 0.5 °C every other day to reach 32 °C in 12 days. The temperature was held constant at 32 °C for 14 days to induce bleaching while minimizing mortality due to sudden tissue necrosis. One coral fragment per colony was sampled at the beginning of the experiment and at subsequent intervals when visible color changes occurred. Due to the mortality of four fragments during the heat phase, only 50 coral samples were included in this analysis.

Corals were held in the main aquarium system with previously described water parameters. A total of eight coral fragments from three colonies of the three species were used (24 fragments per species, 72 fragments in total). The chemical bleaching was conducted in 1-L glass containers filled with 65-µm sieved seawater at 26.5 °C. Corals were incubated with 0.58 mM menthol (stock solution: 1.28 M, 20% w/v menthol in 99% ethanol) for eight hours per day at a light intensity of 200 ± 10 µmol photons m⁻² s⁻¹, stirring (∼2.62 cm/s) in bespoke temperature controlled stirring chambers (Rades et al. 2022). Coral fragments were exposed to three bleaching protocols that differed in the number of incubation days (1, 2, and 4 days) after Bauer et al. (in preparation). Control corals were incubated in menthol-free, sieved seawater. Physiological measurements and imaging were performed 15 days after the last menthol incubation on 68 coral fragments (four fragments died before the end of the experiment).

### RGB data acquisition and processing

To assess the degree of bleaching, color values were extracted from standardized photographs of each coral fragment. Images were taken with a DSLR camera (Nikon D7000 or D7100) equipped with a macro lens (Tamron 90mm macro lens or Nikon AF-S Nikkor 24-70 mm 1:2.8G ED) in an evenly illuminated photo studio (80 × 80 × 80 cm; Life of Photo, Fig. S1). Each fragment was photographed on a black background with the larger side facing the camera, alongside a reference color card for white balance calibration (ColorChecker Passport Photo 2, Calibrite, USA).

Images were processed using Adobe Photoshop (Adobe Inc., USA) and the Meta AI Segment Anything model (Kirillov et al., 2023) for white balance adjustment and background removal. Images were saved in PNG format with a transparent background. Color channel values were extracted from the background-free images using an R script (https://github.com/ErikFerrara/Coral_Color_Index.git) that extracts RGB color channel values from each pixel to generate color histograms. The mean value of all pixels was calculated for each channel and used as a tissue color proxy. Bleaching was scored as a change in tissue color on a scale of 0 to 255, in which higher values correspond to higher bleaching severity. RGB channel values were used to calculate 19 color indices (Table 1) which were previously established as good predictors of total chlorophyll content in plants (Kior et al., 2024).

**Table 1.** – List of the color indices considered in this study, including their equations and sources.

| Index | Name of color parameter | Definition | Reference |
| --- | --- | --- | --- |
| R | Red index | Red (R) channel | (Rigon et al., 2016) |
| G | Green index | Green (G) channel | (Rigon et al., 2016) |
| B | Blue index | Blue (B) channel | (Rigon et al., 2016) |
| r (normalized) | Normalized Red index | $R / (R + G + B)$ | (Rigon et al., 2016) |
| g (normalized) | Normalized Green index | $G / (R + G + B)$ | (Rigon et al., 2016) |
| b (normalized) | Normalized Blue index | $B / (R + G + B)$ | (Rigon et al., 2016) |
| Grayscale | RGB weighted mean | $0.299 \times R + 0.587 \times G + 0.114 \times B$ | (Güneş et al., 2016) |
| R+G | Red-Green sum index |  | (Ali et al., 2012) |
| R+B | Red-Blue sum index |  | (Hu et al., 2010) |
| R-G | Red minus Green index |  | (Chen et al., 2020) |
| R-B | Red minus Blue index |  | (Ali et al., 2012) |
| R/G | Red–Green simple ratio |  | (Sánchez-Sastre et al., 2020) |
| G/R | Green-Red simple ratio |  | (Ali et al., 2012) |
| (R-G)/(R+G) | Normalized Red–Green difference |  | (Kawashima, 1998) |
| (R-B)/(R+B) | Normalized Red–Blue difference |  | (Ali et al., 2012) |
| (G-R)/(G+R) | Green vegetation index |  | (Gitelson et al., 2002) |
| (G-B)/(G+B) | Normalized Green-Blue difference |  | (Kawashima, 1998) |
| R/(G+B) | Simple ratio intensity R-GB |  | (Widjaja Putra & Soni, 2018) |
| (G-R)/(R+G-B) | Visible atmospheric resistance |  | (Widjaja Putra & Soni, 2018) |

## Symbiont cell density and chlorophyll concentration

### Surface Area Calculation

To standardize chlorophyll content and Symbiodiniaceae cell density, the surface area of each coral fragment was determined using either 3D scanning (short– and long-term heat stress experiments) or photogrammetry (menthol bleaching experiment). 3D scanning was performed using an Artec Spider 3D scanner and processed with Artec Studio 16 software (Artec 3D, Luxembourg, version 16.0.8.2). Coral fragments were scanned on a rotating platform inside a uniformly illuminated photobox (Fig. S1). Scans were captured during three complete rotations at different angles—low, middle, and top—to generate high-quality models of each fragment. The 3D models were calculated in Artec Studio 16, with fine registration (features: geometry and texture), global registration (key frame ratio 0.3, search features within mm: 4), sharp fusion (resolution 0.2, fill holes by radius, max. hole radius: 5), and outlier removal. Artifact objects were removed (small objects filter). The processed 3D models were manually trimmed in Artec Studio 16 to calculate the surface area of living coral tissue only.

Photogrammetry modeling was used to generate 3D models of the coral fragments used for the menthol bleaching experiment. Approximately 50 images of each coral fragment were taken before the first menthol incubation. The images were processed using 3DF Zephyr Free software (version 7.003) to create 3D models. Models were trimmed in Artec Studio 16 Professional, and size scaling and surface area calculations were performed using MeshLab (Visual Computing Lab).

### Symbiont cell quantification

For the long-term and short-term heat stress experiments, the tissue was removed using an airbrush (Conrad Electronic, Hirschau, Germany) filled with filtered seawater (FSW). The suspension was homogenized and centrifuged at 3,000 g for 10 min to pellet the symbiont cells. The supernatant was discarded and the pellet was washed and resuspended in 5 mL FSW.

For the menthol bleaching experiment, symbiont cells were separated from the coral skeleton using a 1 M sodium hydroxide (NaOH) solution following Zamoum and Furla (2012). Coral fragments were placed in 50 mL of 1 M NaOH and incubated in a water bath at 36 °C for 60 min and shaken every 15 min. After incubation, the skeleton fragments were removed and the suspension was centrifuged at 3,000 g for 5 min (Laborfuge 400R, Heraeus Instruments). The supernatant was discarded, and the pellet was resuspended in FSW. The washing step was repeated twice, after which the pellet was resuspended in FSW. Symbiodiniaceae cell density was determined using a Thoma cell counting chamber and normalized to tissue surface area.

### Chlorophyll Extraction and Quantification

Chlorophyll content was determined only from the symbiont suspensions obtained by the airbrush method. The suspension was centrifuged and the pellet was washed with DI water three times. The final pellet was resuspended in 10 mL of 100 % acetone and stored in the dark at 0 °C for 24 h to extract chlorophyll pigments. After extraction, samples were centrifuged at 2,000 g for 5 min to remove cellular debris. The absorbance of the supernatants was measured using a spectrophotometer at wavelengths of 630 nm and 663 nm. Chlorophyll *a* and *c2* concentrations were calculated using the equations for dinoflagellates of Jeffrey and Humphrey (1975) and normalized to tissue surface area. Chlorophyll was extracted only from corals exposed to long-term and short-term heat stress. For the menthol bleaching experiment, the use of sodium hydroxide prevented the extraction of chlorophyll.

### Statistical analysis

Statistical analyses were performed using R software (version 4.3.3; R Core Team, 2024). Correlations between color indices and physiological parameters (symbiont cell density and chlorophyll content) were assessed using a linear model. A coefficient of determination (R^2^) and a significance level of p < 0.05 were used to identify and rank the top three coral indices across bleaching conditions for each species. Additionally, root mean square error (RMSE), Akaike information criterion (AIC), and Bayesian information criterion (BIC) were considered to identify the optimal color index across coral species and bleaching conditions.

To accurately predict symbiont densities through color values, we identified the minimum sample size required to train a robust linear model through bootstrap analysis with 1,000 iterations. In each iteration, a random subset of samples, with progressively increasing sample size, was selected to train the linear model. The model was validated on the entire dataset and the Root Mean Square Error (RMSE) between the measured and predicted Symbiodiniaceae cell density was calculated. By evaluating the RMSE across different sample sizes, we identified the smallest sample size required to achieve accurate and reliable predictions. The relative similarity (%) to the RMSE value obtained with the largest sample size (Fig. 4) and the actual RMSE values (Fig. S8) were visualized by a rarefaction-like curve. Thresholds of 90% and 95% similarity were established to determine the minimum sample size required for each treatment condition.

## Results

### Correlation between symbiont cell density and total chlorophyll content

Our initial analysis confirmed a strong positive correlation between symbiont cell density and total chlorophyll content across coral species and bleaching conditions (Fig. 1; R² up to 0.93, p < 0.001). In other words, the chlorophyll content per symbiont cell remained relatively stable across species and bleaching states. Notably, symbiont cell density and chlorophyll in corals exposed to short-term heat stress consistently showed a stronger correlation than those exposed to long-term heat stress (Fig. 1).

**Figure 1.**
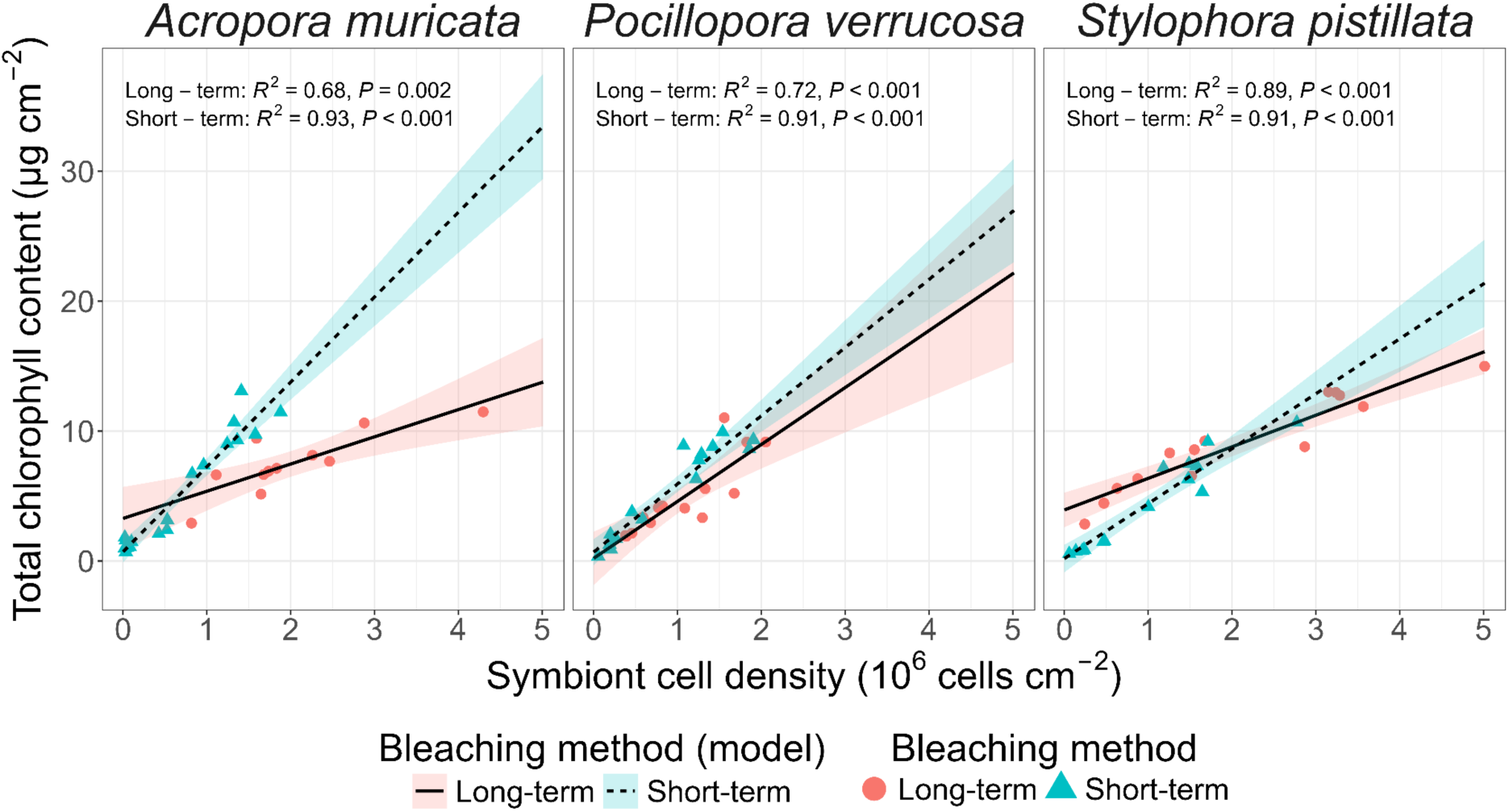
– Relationship between total chlorophyll content and symbiont cell density across bleaching protocols and coral species. The fit for each linear model is shown with the correlation coefficient (R²) and the *p-value*.

### Correlation of color indices with symbiont cell density

In our assessment of RGB-derived color indices for predicting symbiont cell density, we found that indices successfully used to predict chlorophyll content in plants differed in their ability to predict symbiont densities in corals (Fig. 2). Indices such as the Red index (R), the Red-Green sum (R+G) index, and the Grayscale index correlated strongly with symbiont density across all experimental conditions.

**Figure 2.**
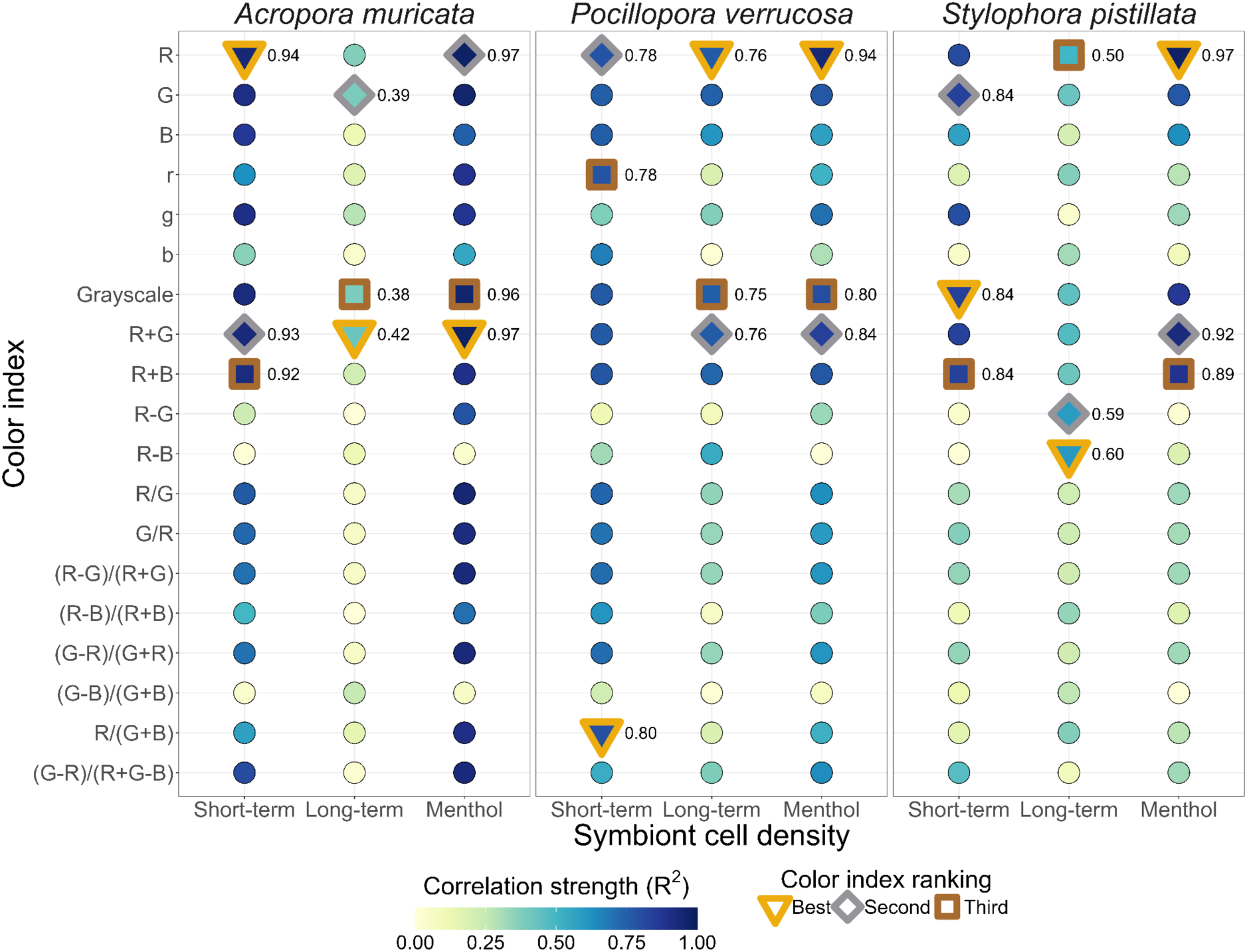
– Coefficients of determination (R²) of different color indices against symbiont cell density across species and bleaching treatments. Linear regression models were fitted on raw data separately for each bleaching method. The color of each point represents the R² value, illustrating the strength of the correlation: the higher the correlation, the darker the color. Points with a thick border highlight the top three R² values, indicating the best color indices to use as a proxy for symbiont cell density estimation.

The Red index proved to be a particularly robust predictor of symbiont density across species and bleaching conditions, with the highest R² in four out of nine experimental conditions (Fig. 2). In coral species bleached with menthol, the R² value of this index was highest at up to 0.97, indicating a strong linear correlation with symbiont cell density. Overall bleaching treatments, the Red index had an average R² of 0.82 and 0.76 in *P. verrucosa* and *S. pistillata*, respectively. In *A. muricata*, the Red-Green sum index (R+G) had a slightly higher overall R² value (0.77) than the Red index (0.76), but the differences were small. Following the Red index, the Red-Green sum and Grayscale indices also demonstrated high R² values, making them additional candidate proxies.

The Red index consistently reduced the spread of data points along the x-axis for severely bleached coral fragments (Fig. 3). This was particularly pronounced in the short-term and menthol bleaching treatments where these data points were tightly clustered, resulting in a good linear model fit. In addition to R², statistical metrics such as *p*-values, Akaike Information Criterion (AIC), and Bayesian Information Criterion (BIC) consistently ranked the Red index best to describe the linear correlation with symbiont cell density across bleaching conditions and species (Table 2).

**Figure 3.**
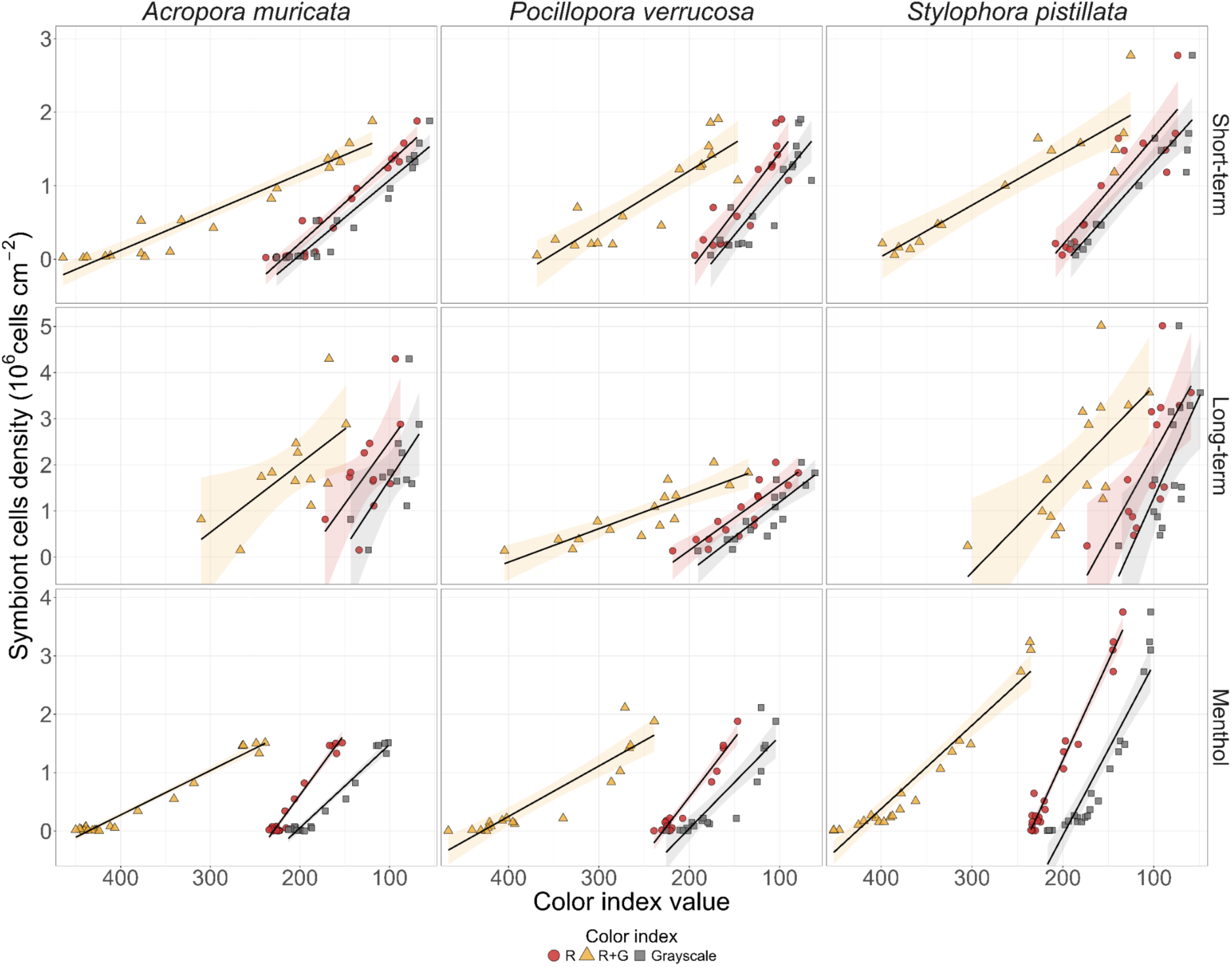
– The three best color indices (Red, Red-Green sum, and Grayscale) to predict symbiont cell density across coral bleaching methods. Linear regression models were fitted separately for each color index on raw data. The coefficient of determination (R^2^) and *p-*values were obtained from raw data after removing highly influential points (based on Cook’s distance Fig. S9-S11).

**Figure 4.**
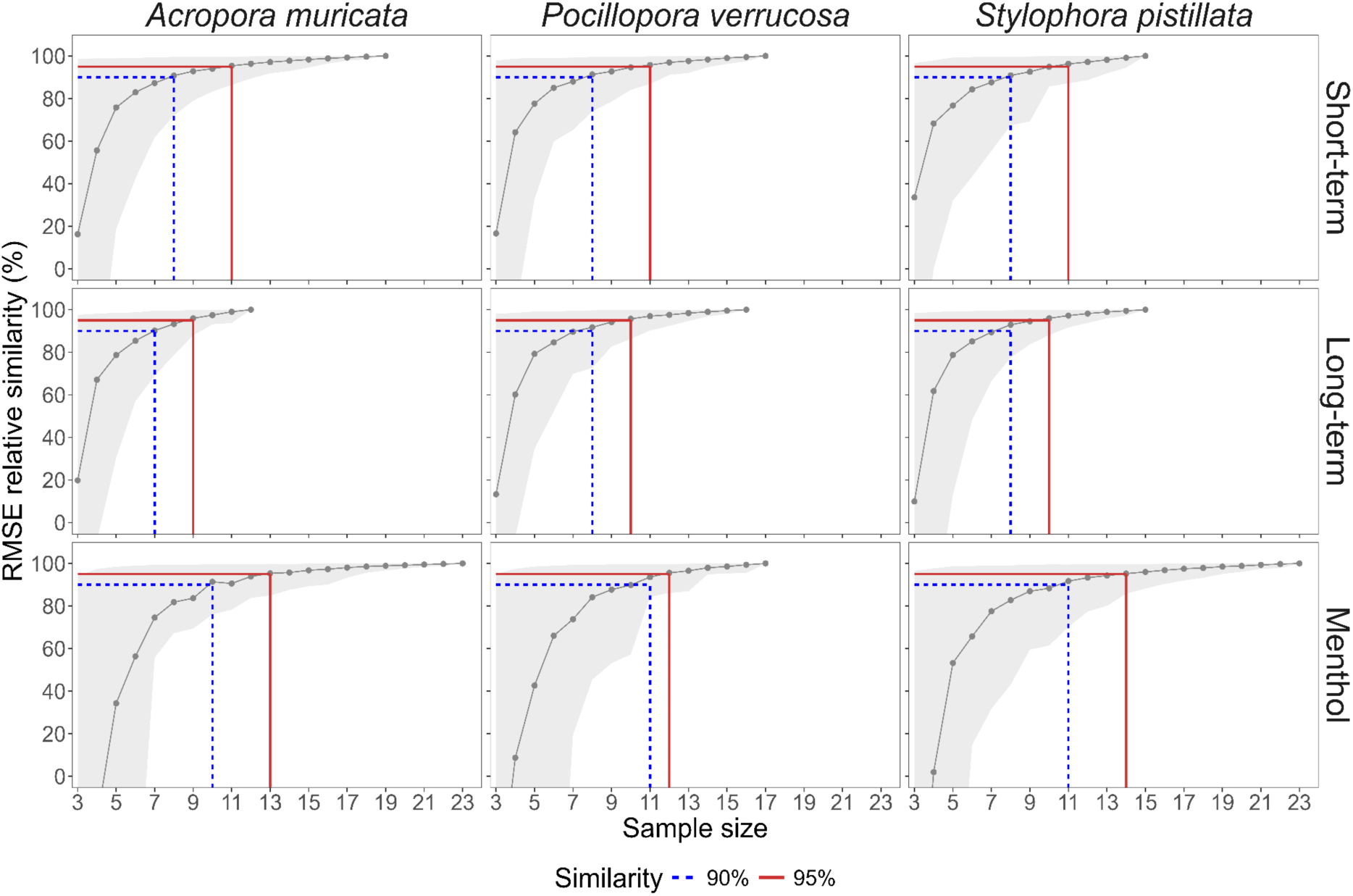
– Determining the minimum sample size for training a linear model to predict symbiont cell density from coral tissue color. The similarity of the RMSE metric between models fit on an increasing sample size and the model fit on all observations (mean values calculated on a bootstrap of 1,000, shaded area = 90 % CI). Thresholds of 90 and 95 % similarity between training and full data models are highlighted to indicate the sample size required for consistent model performance.

**Table 2.**
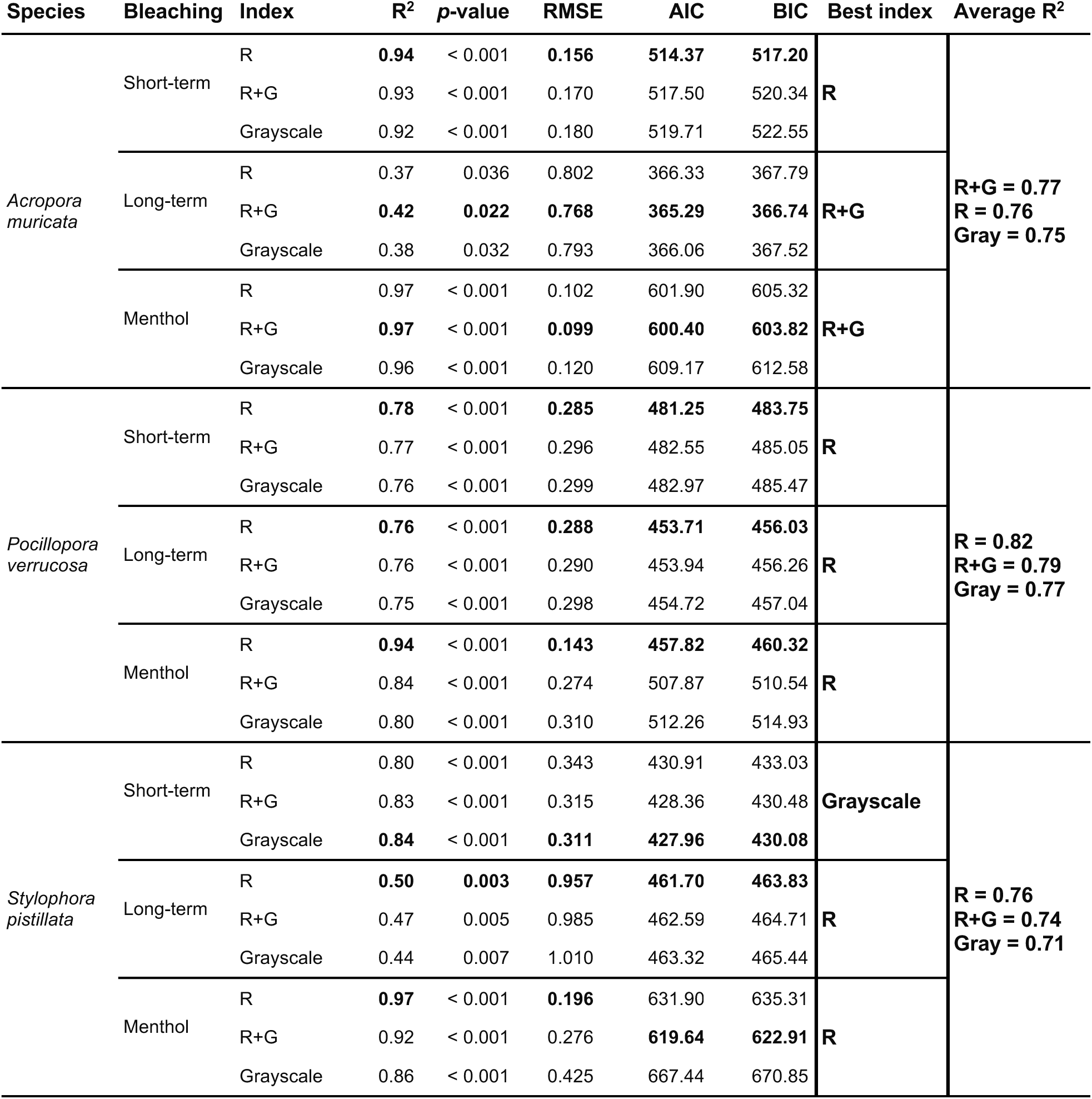
– Statistical overview of the best color indices as proxies for symbiont cell density across coral species and bleaching conditions. Indices were ranked based on the R^2^, the *p*-value, the root mean square error (RMSE), the Akaike information criterion (AIC), and the Bayesian information criterion (BIC) of models predicting symbiont density. The best scores for each criterion were highlighted in bold.

When exposed to long-term heat stress, the correlation of all color indices with symbiont density was lower than after short-term and menthol bleaching across all coral species. For instance, in *A. muricata*, the highest R² value was 0.42 for the Red-Green sum index, which was lower than in the other bleaching treatments (0.93 and 0.97). Other indices generally performed less well and were therefore excluded from further analysis.

### Correlation of color indices with total chlorophyll content

Extending our analysis predicting total chlorophyll content from coral tissue colors, we found that it was best described by the same indices as symbiont cell density, although small differences occurred (Fig. S3). Specifically, the Red index was the best index to describe the linear correlation with total chlorophyll content across all bleaching conditions and species and the Red-Green sum (R+G) index performed equally well (Fig. S4, Table S1). Additionally, the Red-Blue sum (R+B) index performed better as a chlorophyll proxy than the Grayscale index (Fig. S4, Table S1). The correlation strength between the Red index and chlorophyll content was slightly better (or equal) than with symbiont density in most of the bleaching conditions.

### Colorful bleaching and endolithic algae

We found that the occurrence of colorful bleaching in our sample set did not strongly affect the correlation between color values and symbiont cell densities and chlorophyll content (Figs. S5, S6). However, a high amount of endolithic algae within the coral skeletons negatively affected the color values in a few cases, as it was observed to be outside the standard error interval of the linear model (Fig. S5, *A. muricata* exposed to long-term heat stress).

### Determining minimum sample size requirements for robust linear regression

We assessed the minimum sample size necessary for training a reliable linear model to predict symbiont cell density and chlorophyll content from Red index values. The similarity of the root mean square error (RMSE) of models trained on data subsets, relative to the RMSE of a model trained on the entire dataset increased with sample size, suggesting a significant improvement in the predictive accuracy with higher sample numbers (Fig. 4). Based on these analyses 7-9 samples were needed to train a model with 90 % accuracy and 9-14 samples to reach 95 % accuracy (Fig. 4).

## Discussion

We demonstrated that the Red index is a robust predictor of coral health, consistently showing a strong linear correlation with Symbiodiniaceae cell density and total chlorophyll content in *Acropora muricata*, *Pocillopora verrucosa*, and *Stylophora pistillata*, exposed to acute short-term heat stress, chronic long-term heat stress, and menthol-induced bleaching. Furthermore, the strong correlation between the Red index and coral health parameters remains stable even in the presence of colorful bleaching or endolithic algae, confirming that image analysis based on the Red index is a rapid, reliable, and non-invasive tool for assessing coral health in both large-scale monitoring initiatives and laboratory experiments.

### Color changes scale linearly with symbiont cell density

Our systematic assessment demonstrated the Red index to be linearly correlated with two key physiological parameters commonly used in coral health assessments: symbiont cell density and total chlorophyll content (Grottoli et al., 2021; Jones, 1997). Such linearity implies that changes in Red index values can be directly interpreted as proportional changes in symbiont density or total chlorophyll content regardless of coral bleaching severity. In contrast, the other indices resulted in an overall lower correlation strength driven by large color variations among severely bleached coral fragments and smaller color variations among healthy fragments in comparison to the same change in symbiont cell density. We propose the Red index as the best-performing proxy for assessing coral health through image analysis, as it consistently showed the highest correlation across coral species and bleaching conditions.

Our findings align with other studies in plant physiology, where the Red index is commonly used as a proxy for chlorophyll content in leaves (R^2^ up to 0.94, Hu et al., 2010; Riccardi et al., 2014; Rigon et al., 2016). Image analysis has become a common proxy for traditional bleaching metrics in several coral studies (Evensen et al., 2022; Reichert et al., 2021; Teague et al., 2022; Voolstra et al., 2020), though the Red index has only been confirmed as a reliable proxy for total chlorophyll content for a small number of coral species. For instance, Winter et al. (2009) demonstrated the correlation between the Red index and chlorophyll content (R^2^ 0.82) in *Stylophora pistillata*. Additionally, although changes in symbiont density have previously been correlated with RGB-based color score matrices (Edmunds et al., 2003; Strand et al., 2024) or categorical color scales (Bahr et al., 2020; Siebeck et al., 2006), our systematic assessment proved for the first time that the Red index is also strongly correlated with symbiont cell density. The highest linearity of the Red index across coral species exposed to different bleaching conditions here confirms its use as a proxy for chlorophyll content and allows for fast and straightforward assessments of symbiont cell density.

The strength of the correlation between coral color and symbiont cell density and chlorophyll content differed. Particularly, in some of our study cases, the Red index was more strongly correlated with chlorophyll content than symbiont density. It is known that during coral bleaching, the decrease in the total chlorophyll content in corals is mainly due to a loss of Symbiodinaceae cells, rather than the loss of chlorophyll content per symbiont cell. However, in some cases, chlorophyll is degraded through photo-oxidative reactions, resulting in a loss of pigmentation without a decrease in symbiont cell density (Venn et al., 2006). Accordingly, the slightly weaker correlation between the Red index and symbiont cell density than with chlorophyll content in our work may indicate that the coral host retained a portion of symbiont cells despite the loss of photosynthetic pigments. This observation contrasts with Siebeck et al. (2006), who reported that the color score measured with the Coral Health Chart correlated more strongly with symbiont density than with chlorophyll content (R² = 0.63 and 0.36, respectively). These contrasts may be explained by differences in experimental conditions, coral and symbiont species, or their geographical origin, which are known to influence the visual appearance of coral colonies (Edmunds et al., 2003; Fitt et al., 2000; Hackerott et al., 2023, 2024; Strand et al., 2024). This variability in coral visual appearance also necessitates region-specific Coral Health Charts to ensure reliable coral health assessments based on color matrics (Bahr et al., 2020; Siebeck et al., 2006). Similarly, the method implemented here requires careful consideration based on the experimental conditions. Although the Red index shows a strong linear correlation with coral health parameters within each experimental setup, comparing absolute color values or their changes across different experiments is less reliable as indicated by the different regression slopes.

We have demonstrated that training linear models on Red index values allows reliable prediction of relational changes in symbiont densities and chlorophyll content. However, predicting exact symbiont densities or chlorophyll content requires the training of a bespoke linear model for each sample set. To train a high-performance model, comparable to those trained on the full dataset in our work (similarity >95%), we suggest using 10-12 samples. The number of samples may possibly be reduced if samples from different stages of bleaching severity are included for model training. We have provided a ready-to-use R script to extract color values from trimmed coral images to streamline the data analysis workflow.

### Challenges and limitations

Overall, we have demonstrated that assessing coral health based on the Red index is a rapid and robust alternative to traditional invasive bleaching metrics, which outperforms all other indices tested in this work. Nonetheless, researchers should be aware of the limitations inherent to this method, such as condition-specific differences affecting color measurements. Given that a bleaching response is the endpoint of several biological mechanisms and physiological processes (Helgoe et al., 2024), it is reasonable that differences in the bleaching trigger may affect the visual appearance of corals. Such stimuli can induce physiological acclimatization or changes in pigmentation that are not directly related to Symbiodiniaceae loss (Bollati et al., 2020; Fitt et al., 2000; Shick & Dunlap, 2002). For instance, prolonged exposure to high temperatures may promote the proliferation of endolithic algae within the coral skeleton, potentially affecting the color analyses of bleached corals. In our experiment, the presence of endolithic algae may partially explain the lower performance of the Red index in the long-term heat exposure experiment, where the proliferation of endolithic algae caused additional pigmentation in some of the severely bleached corals, thereby altering color index values.

Another important phenomenon that may interfere with color analysis is the so-called colorful bleaching. It is caused by the upregulation of host-derived fluorescent pigments and chromoproteins in response to stress (Bollati et al., 2020), and the accumulation of such non-photosynthetic pigments leads to tissue coloration (e.g., violet, blue, or green) that masks the loss of symbiont cells. Interestingly, our analysis suggests that colorful bleaching did not affect Red index values when it occurred.

Particularly, in menthol-bleached corals, all changes in Red index values closely correspond to changes in symbiont cell density regardless of the visible accumulation of non-photosynthetic pigments.

### Recommendations for future applications

We emphasize that high-quality data acquisition is crucial to maximize the reliability of image-based analyses. Images should be captured in a uniformly illuminated environment to reduce shadows and avoid overexposure. Under laboratory conditions, this can be achieved using an evenly illuminated photo box studio (Figs. S1, S2). When pictures are not taken underwater, it is important to minimize excess water on coral surfaces to prevent “sun glint” contamination, which can distort colors and brightness. In the field, using artificial lighting setups and a color reference card may help reduce environmental variability (Winters et al., 2009).

The investigation of RGB-based indices commonly used in plant studies has revealed that indices such as the Red-Green and Red-Blue sum are also strongly correlated with coral physiological parameters. Interestingly, in terrestrial botany, RGB-based indices are also used as a proxy for other parameters, including plant nitrogen concentration (Lu et al., 2021; Zheng et al., 2018) and estimation of plant biomass (Schirrmann et al., 2016), which incentivizes the exploration of such links with regards to coral color. Further research employing advanced imaging techniques such as hyperspectral imaging, which captures a wider range of wavelengths, could enhance the performance of these indices or lead to the identification of novel indices with higher performance. Moreover, pairing image analysis with other physiological or molecular parameters may help identify indices that predict additional aspects of coral health, such as thermal tolerance or stress response capacity (Hackerott et al., 2024). Moreover, integrating multiple data sources may improve the robustness of health assessments and contribute to a more comprehensive understanding of coral responses to environmental stressors.

### Conclusions

Our research has shown that the Red index is a strong and accurate indicator of symbiotic cell density and chlorophyll content for all coral species and bleaching conditions. The strong linear relationship between these metrics simplifies the interpretation of color value changes, allowing rapid and non-invasive assessment of coral health and stress responses based on tissue coloration. This is particularly important in the face of the increasing threats posed by climate change to coral reef ecosystems, where a reliable tool to rapidly estimate coral health status can fasten the development of effective strategies to mitigate stress in corals. Future research using advanced imaging techniques and additional physiological metrics can further enhance coral health monitoring and conservation efforts.

## Acknowledgements

The authors would like to express their gratitude to the staff of the *Ocean2100* facilities of Justus Liebig University Giessen for their support in animal care and the maintenance of research facilities. Special thanks are also extended to the Marine Holobiomics Lab for their assistance and daily support, and Wyatt Million for input during the initial phase of the project.

## Author contributions

E.F.F., J.R., and M.Z. conceived and designed the study. E.F.F., L.B., J.R., G.P., A.A., and the XMB 2023 student group collected data. E.F.F. analyzed and curated data. E.F.F. and M.Z. wrote and revised the manuscript. M.Z. provided research materials and logistics. All authors read and approved the manuscript.

## Availability of data and materials

Data, analyses, and visualizations are presented in the Supplementary materials. All original data and analysis code are accessible via the GitHub repository under the accession link: https://github.com/ErikFerrara/Coral_Color_Index.git

## Declarations

The authors declare that they have no competing interests.

## Supplementary material

**Figure S1.**
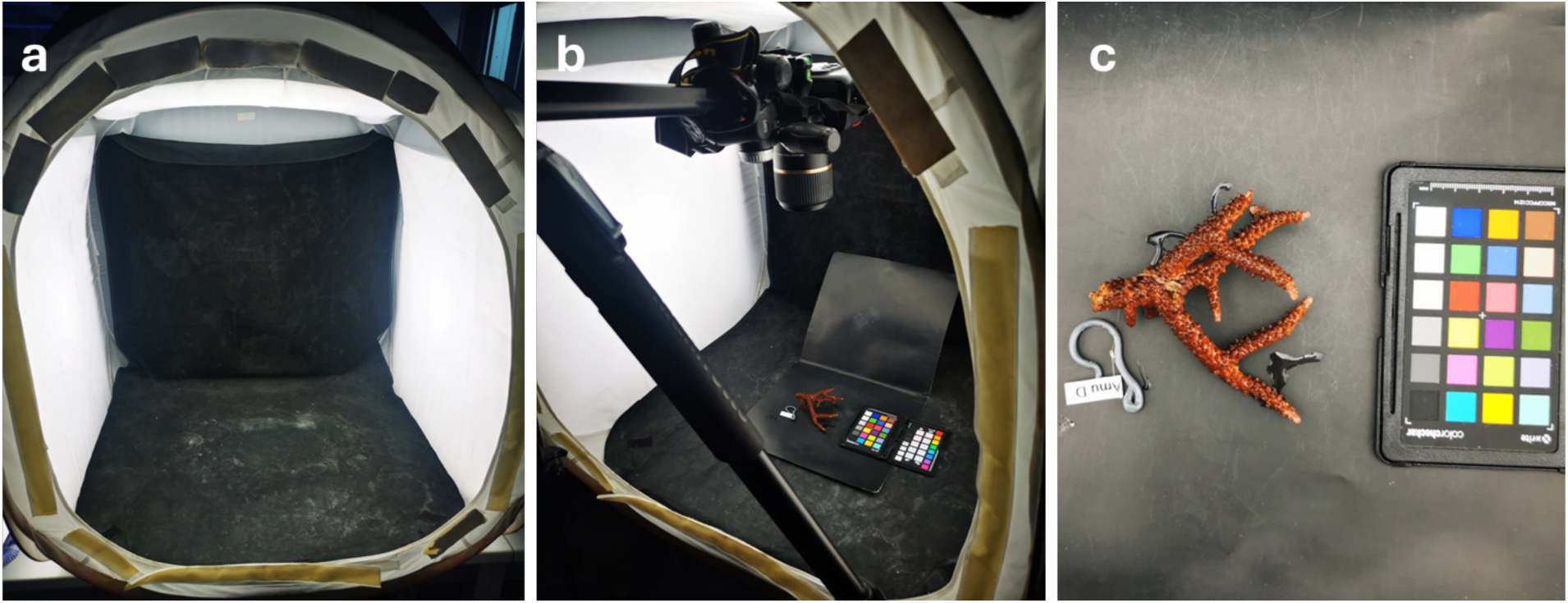
– Setup for standardized photography of the corals. A uniformly illuminated photo box was used to capture images of the corals in all experiments (a). The camera was positioned directly above the corals to minimize any shading caused by the direction of the light sources (b). A color palette was also placed next to the corals for reference (c). Avoiding strong external lights (e.g., sunlight or colorful artificial lights) that might affect the image quality is recommended.

**Figure S2.**
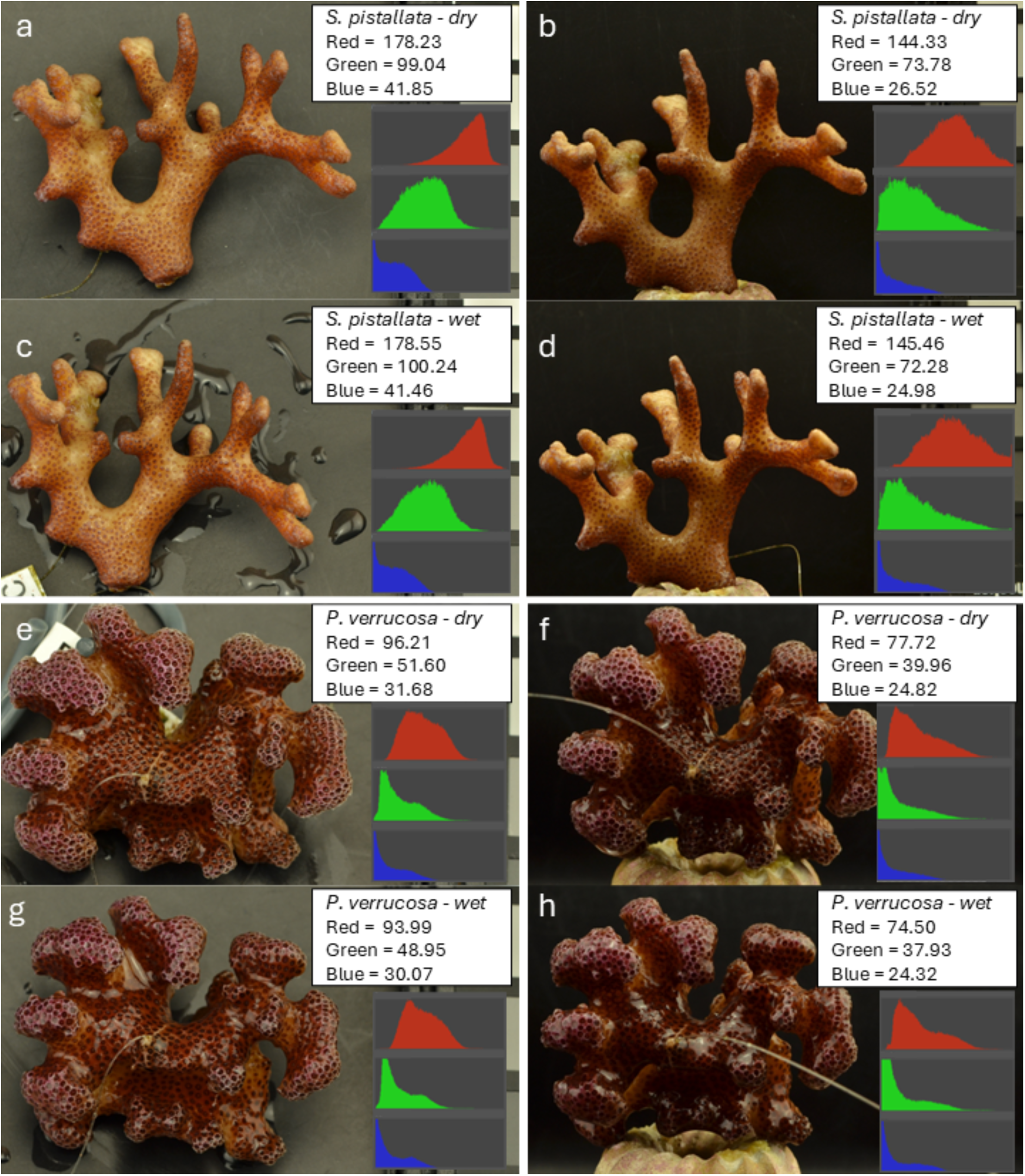
– Effect of different light angles on RGB colors. Images of the same coral were captured in the same evenly illuminated photo box but facing the light from different angles. In A, C, E, and F the captured coral side faces the light directly; in B, D, F, and H the light comes from above, increasing the shaded areas. The “sun glint” effect is stronger in corals that retain more water, which depends on the coral species morphology. The values shown in the images and histograms (red, green, and blue channels, respectively) were extracted from the coral with the background removed.

**Figure S3.**
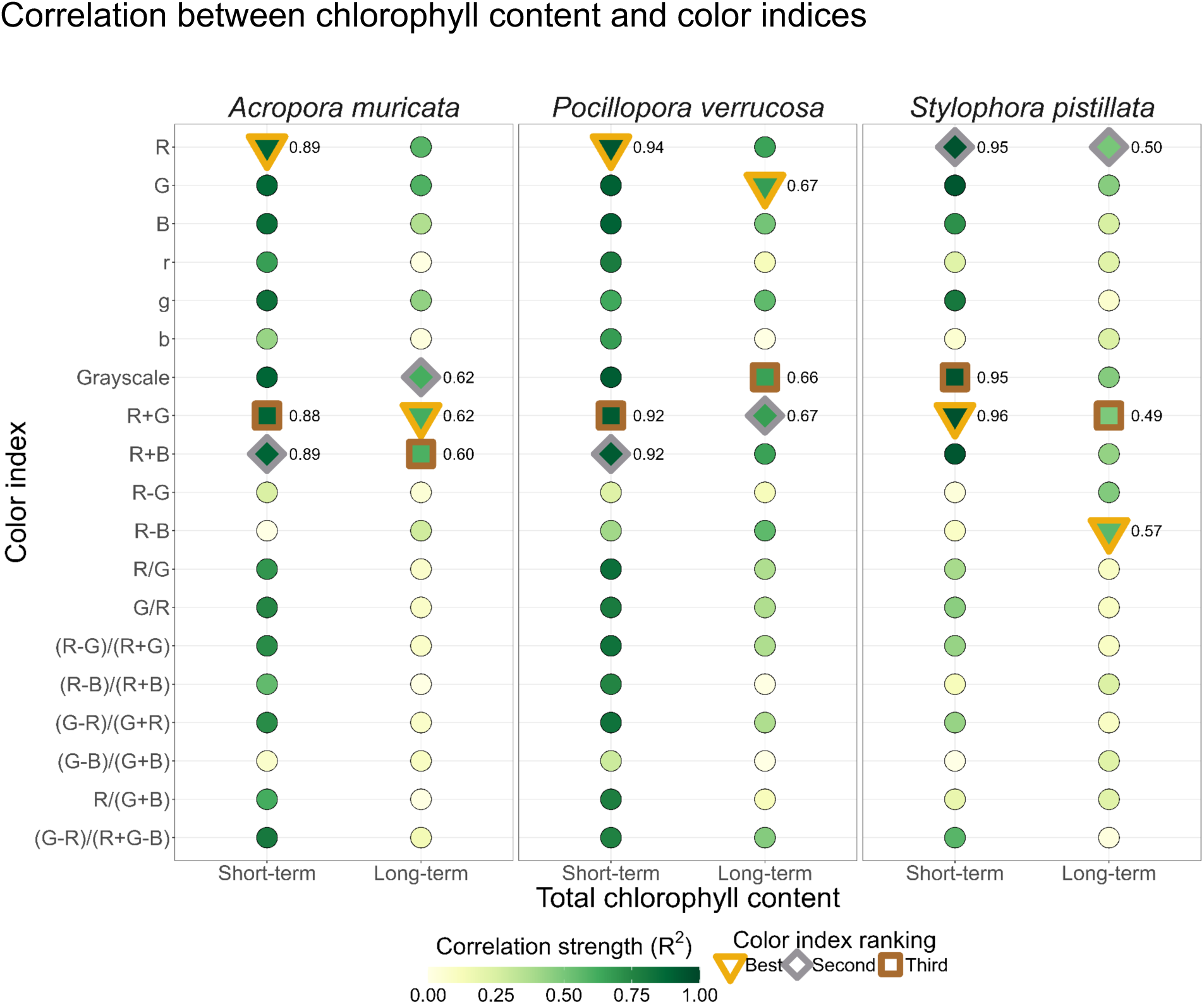
– Coefficients of determination (R²) of different color indices against chlorophyll content across species and bleaching treatments. Linear regression models were fitted on raw data separately for each bleaching method. The color of each point represents the R² value, illustrating the strength of the correlation: the higher the correlation, the darker the color. Points with a thick border highlight the top three R² values, indicating the best color indices to use as a proxy for chlorophyll content estimation.

**Figure S4.**
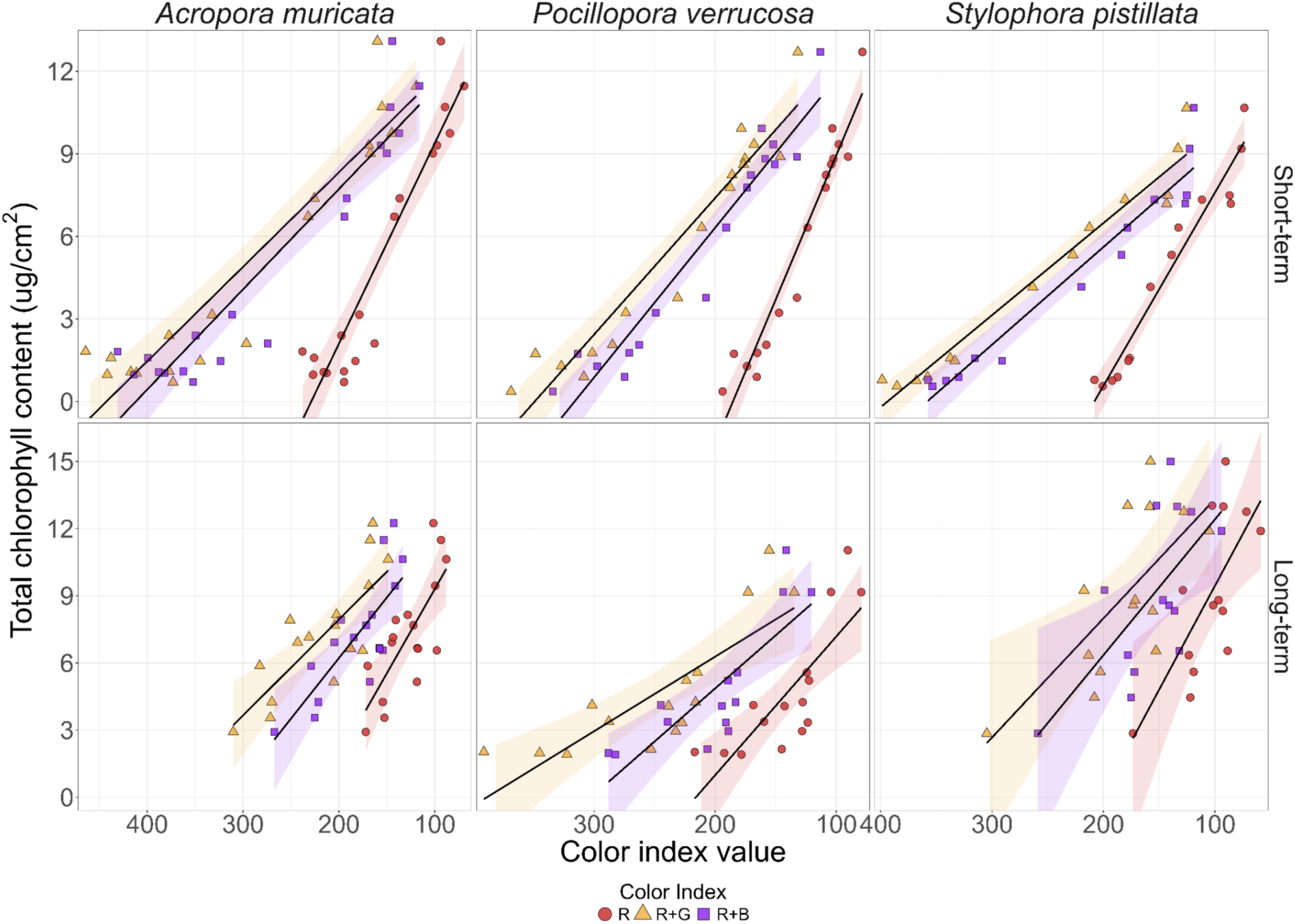
– The three best color indices (Red, Red-Green sum, and Red-Blue sum) to predict chlorophyll content across coral bleaching methods. Linear regression models were fitted separately for each color index on raw data.

**Table S1.** – Statistical overview of the best color indices as proxies for total chlorophyll content across coral species and bleaching conditions. Indices were ranked based on the R^2^, the p-value, the root mean square error (RMSE), the Akaike information criterion (AIC), and the Bayesian information criterion (BIC) of models predicting total chlorophyll content. The best scores for each criterion were highlighted in bold.

| Species | Bleaching | Index | R<sup>2</sup> | p-value | RMSE | AIC | BIC | Best | Average R<sup>2</sup> |
| --- | --- | --- | --- | --- | --- | --- | --- | --- | --- |
| Acropora muricata | Short-term | R | 0.89 | < 0.001 | 1.4 | 72.70296 | 75.53628 | R | R+G = 0.75<br>R+B = 0.74<br>R = 0.73 |
|  |  | R+G | 0.88 | < 0.001 | 1.434 | 73.60596 | 76.43928 |  |  |
|  |  | R+B | 0.89 | < 0.001 | 1.415 | 73.11418 | 75.9475 |  |  |
|  | Long-term | R | 0.58 | 0.0003 | 1.649 | 71.25484 | 73.75448 | R+G |  |
|  |  | R+G | 0.62 | 0.0001 | 1.571 | 69.59441 | 72.09405 |  |  |
|  |  | R+B | 0.6 | 0.0002 | 1.6 | 70.22562 | 72.72526 |  |  |
| Pocillopora verrucosa | Short-term | R | 0.94 | < 0.001 | 0.955 | 52.68238 | 55.18202 | R | R = 0.80<br>R+G = 0.80<br>R+B = 0.79 |
|  |  | R+G | 0.92 | < 0.001 | 1.081 | 56.88013 | 59.37977 |  |  |
|  |  | R+B | 0.92 | < 0.001 | 1.06 | 56.21005 | 58.70969 |  |  |
|  | Long-term | R | 0.65 | 0.0003 | 1.658 | 63.73546 | 65.85961 | R+G |  |
|  |  | R+G | 0.67 | 0.0002 | 1.619 | 63.02096 | 65.14512 |  |  |
|  |  | R+B | 0.66 | 0.0004 | 1.643 | 59.63553 | 61.5527 |  |  |
| Stylophora pistillata | Short-term | R | 0.95 | < 0.001 | 0.778 | 38.694 | 40.61117 | R+G | R = 0.72<br>R+G = 0.72<br>R+B = 0.69 |
|  |  | R+G | 0.96 | < 0.001 | 0.708 | 36.07598 | 37.99315 |  |  |
|  |  | R+B | 0.94 | < 0.001 | 0.845 | 41.02061 | 42.93779 |  |  |
|  | Long-term | R | 0.5 | 0.005 | 2.493 | 71.31146 | 73.22863 | R |  |
|  |  | R+G | 0.49 | 0.005 | 2.522 | 71.63304 | 73.55021 |  |  |
|  |  | R+B | 0.44 | 0.01 | 2.653 | 73.0461 | 74.96327 |  |  |

**Figure S5.**
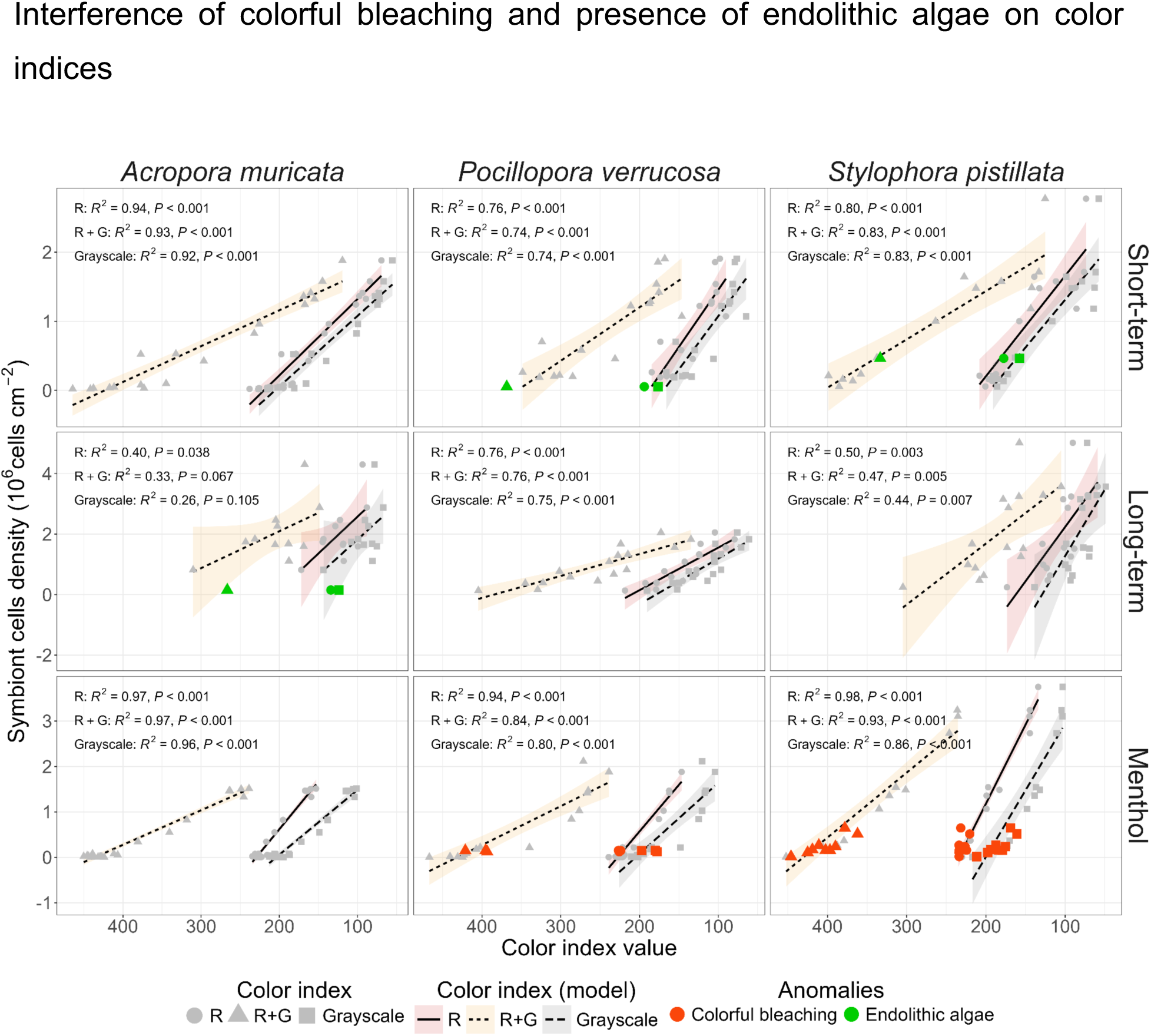
– Distribution of coral samples with color anomalies (endolithic algae and colorful bleaching). Visual distribution of coral samples exhibiting colorful bleaching (red) or endolithic algae (green) and the effect of their exclusion on the linear regression performances.

**Figure S6.**
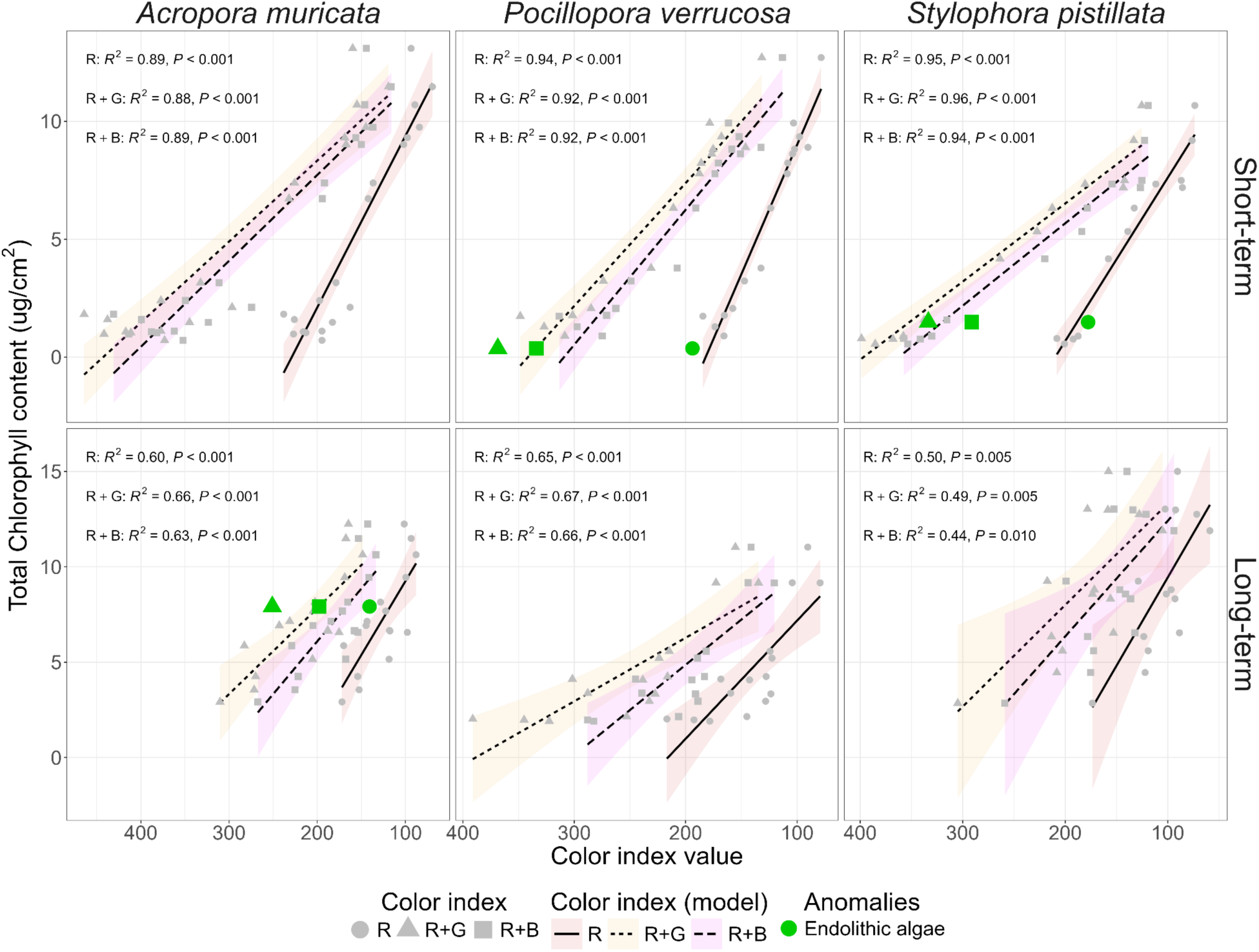
– Visual distribution of coral samples exhibiting endolithic algae (green) and the effect of their exclusion on the linear regression performances.

**Figure S7.**
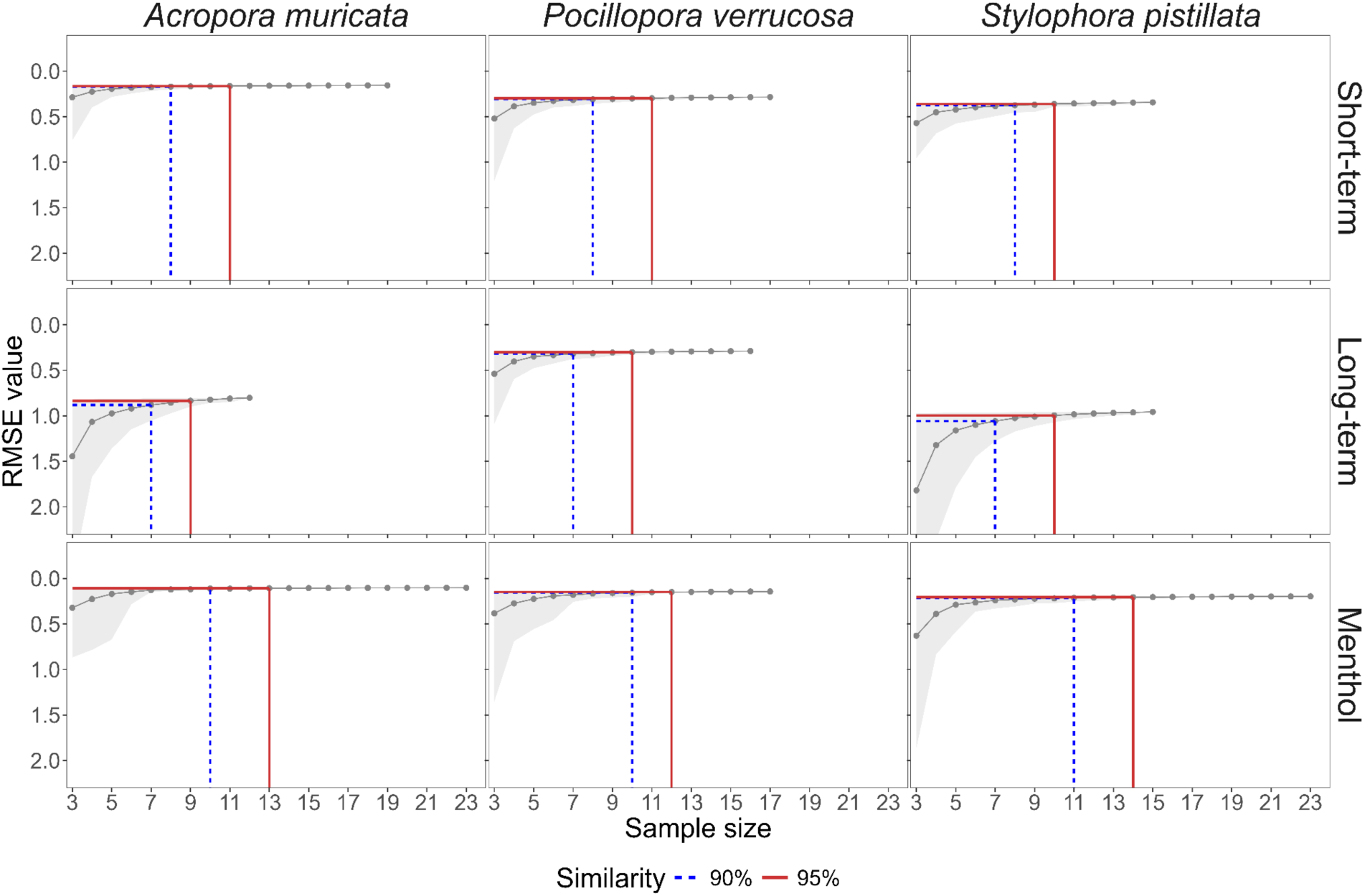
– Determining the minimum sample size for training a linear model. The graph shows the RMSE values of models trained with increasing sample size (mean values calculated on a bootstrap of 1,000). Thresholds of 90 and 95 % similarity are highlighted to indicate the smallest sample size required for consistent model performance comparable to the model trained on the full dataset. The RMSE values represent the precision, expressed in 10^6^ symbiont cells per cm², of the model in predicting symbiont density.

**Figure S8.**
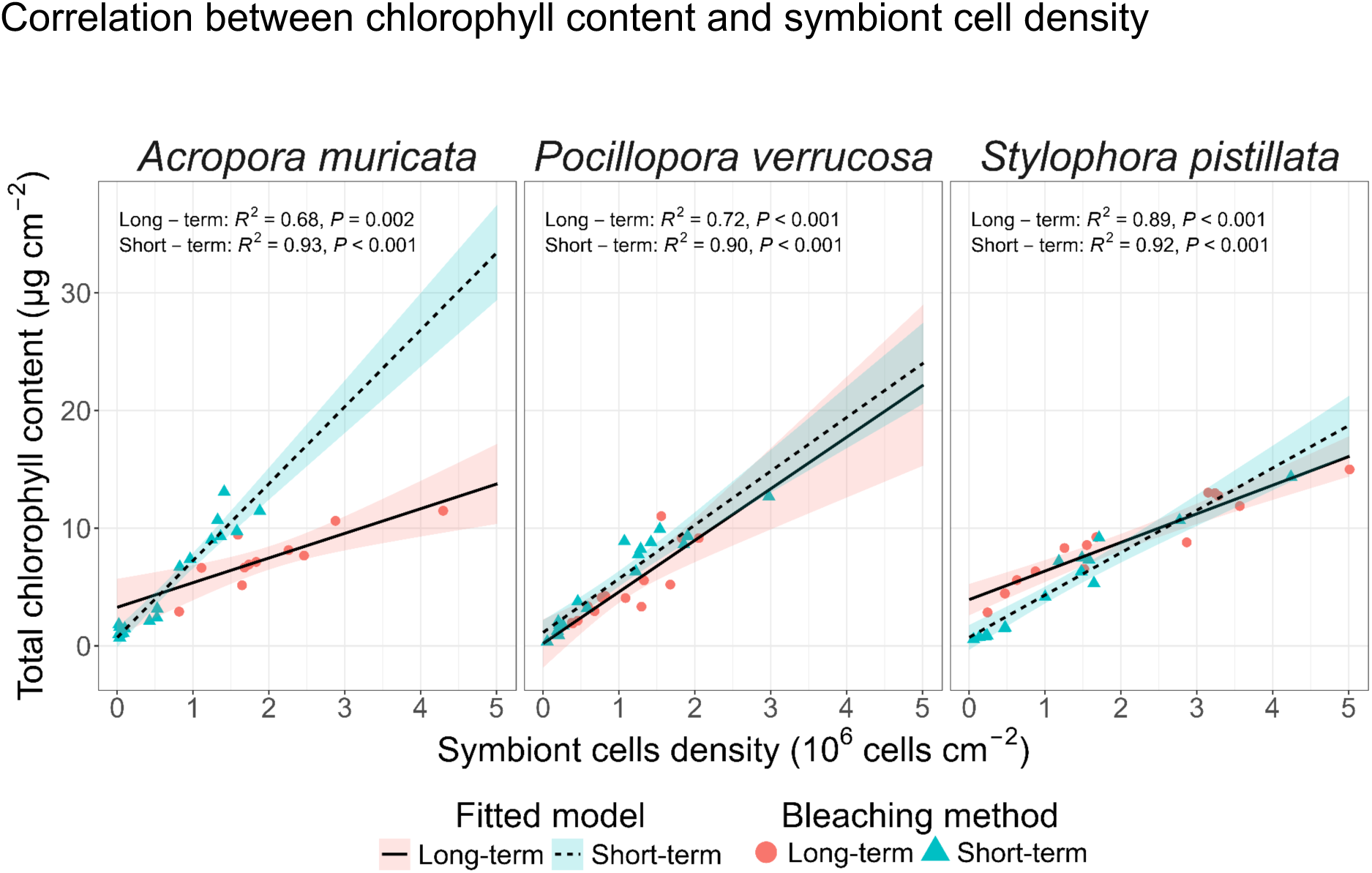
– Relationship between total chlorophyll content and symbiont cell density across bleaching protocols and coral species (with outliers, but excluding the menthol bleaching experiment due to the lack of data). The statistical fit for each linear model is annotated (correlation coefficient (R²) and the *p-value*) and the intercept of the linear model with the y-axis allows for quick assessment of the strength and significance of the relationships observed.

**Figure S9.**
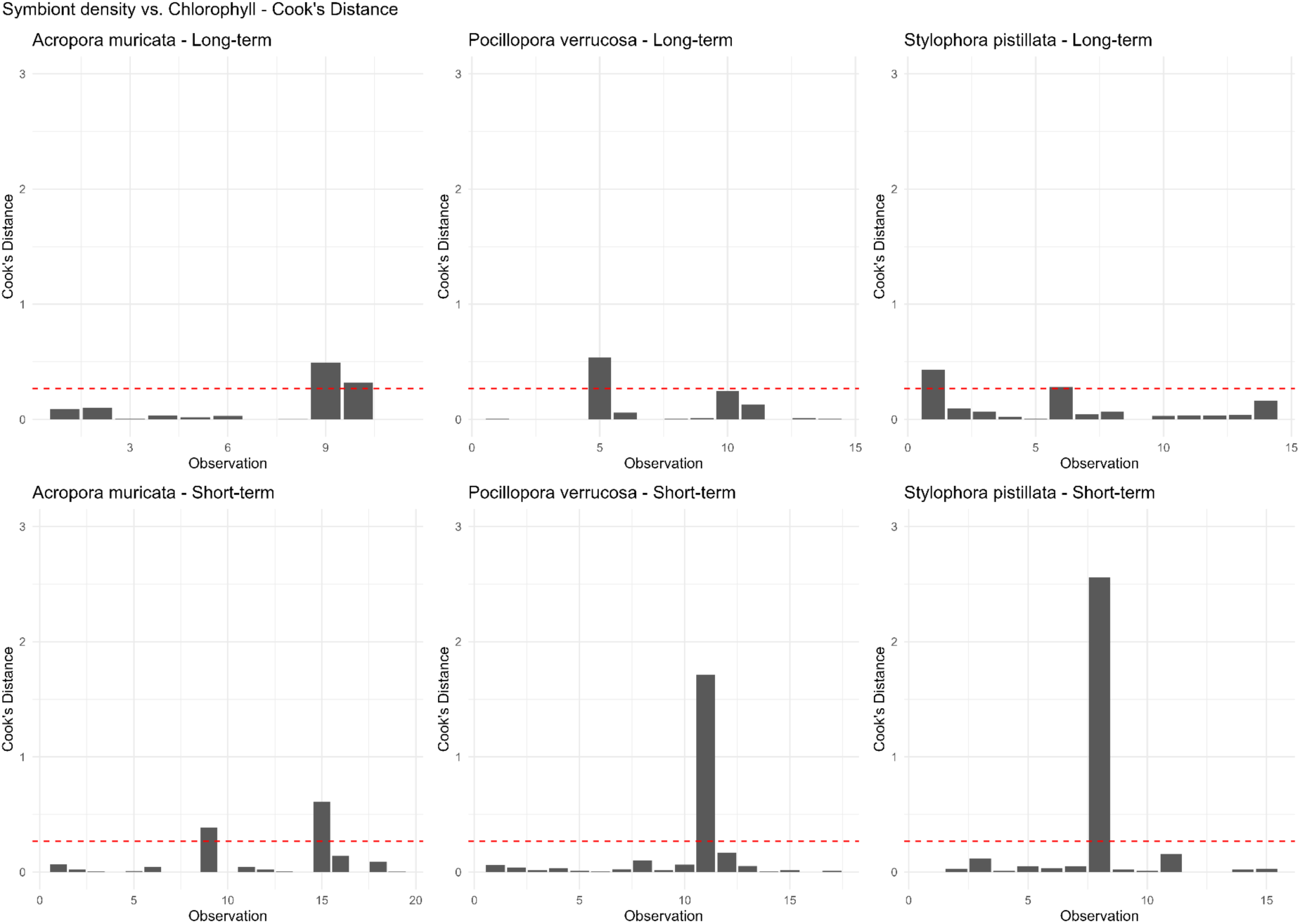
– Samples cook’s distance (D_i_) in the regression model between total chlorophyll content and symbiont cell density across bleaching protocols and coral species. The plot above represents Cook’s distance for each sample in a linear regression model that examines the relationship between symbiont cell density (independent variable) and chlorophyll content (dependent variable). D_i_ is a measure used to identify influential observations (outliers) in the dataset that may disproportionately affect the regression results. Values above the dashed line need to be investigated and eventually removed.

**Figure S10.**
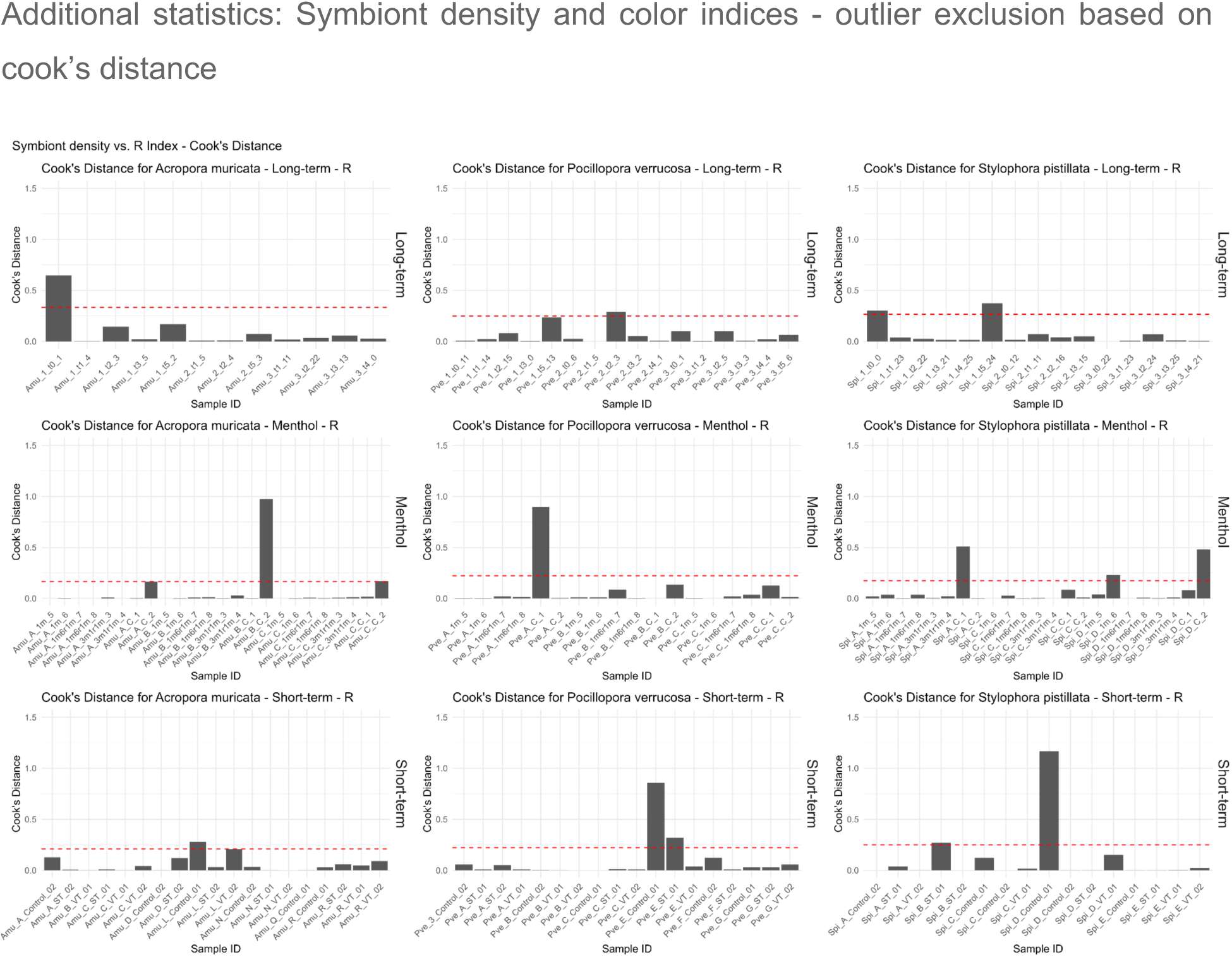
– Samples cook’s distance (D_i_) in the regression model between symbiont cell density and Red index (R) across bleaching protocols and coral species. D_i_ is a measure used to identify influential observations (outliers) in the dataset that may disproportionately affect the regression results. Only values with a D_i_ > 0.7 were removed.

**Figure S11.**
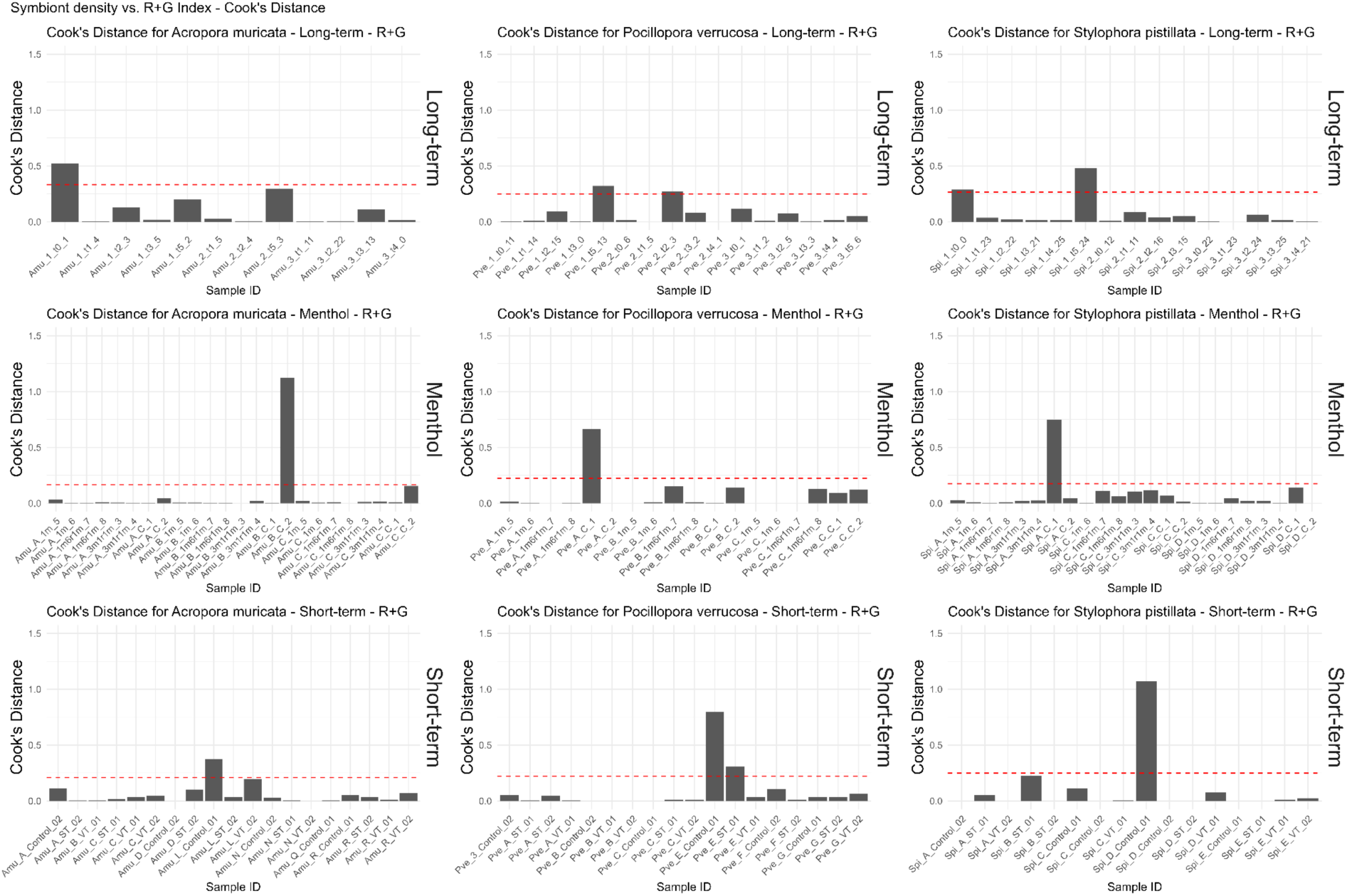
– Samples cook’s distance (D_i_) in the regression model between symbiont cell density and red-green sum index (R+G) across bleaching protocols and coral species. D_i_ is a measure used to identify influential observations (outliers) in the dataset that may disproportionately affect the regression results. Only values with a D_i_ > 0.7 were removed.

**Figure S12.**
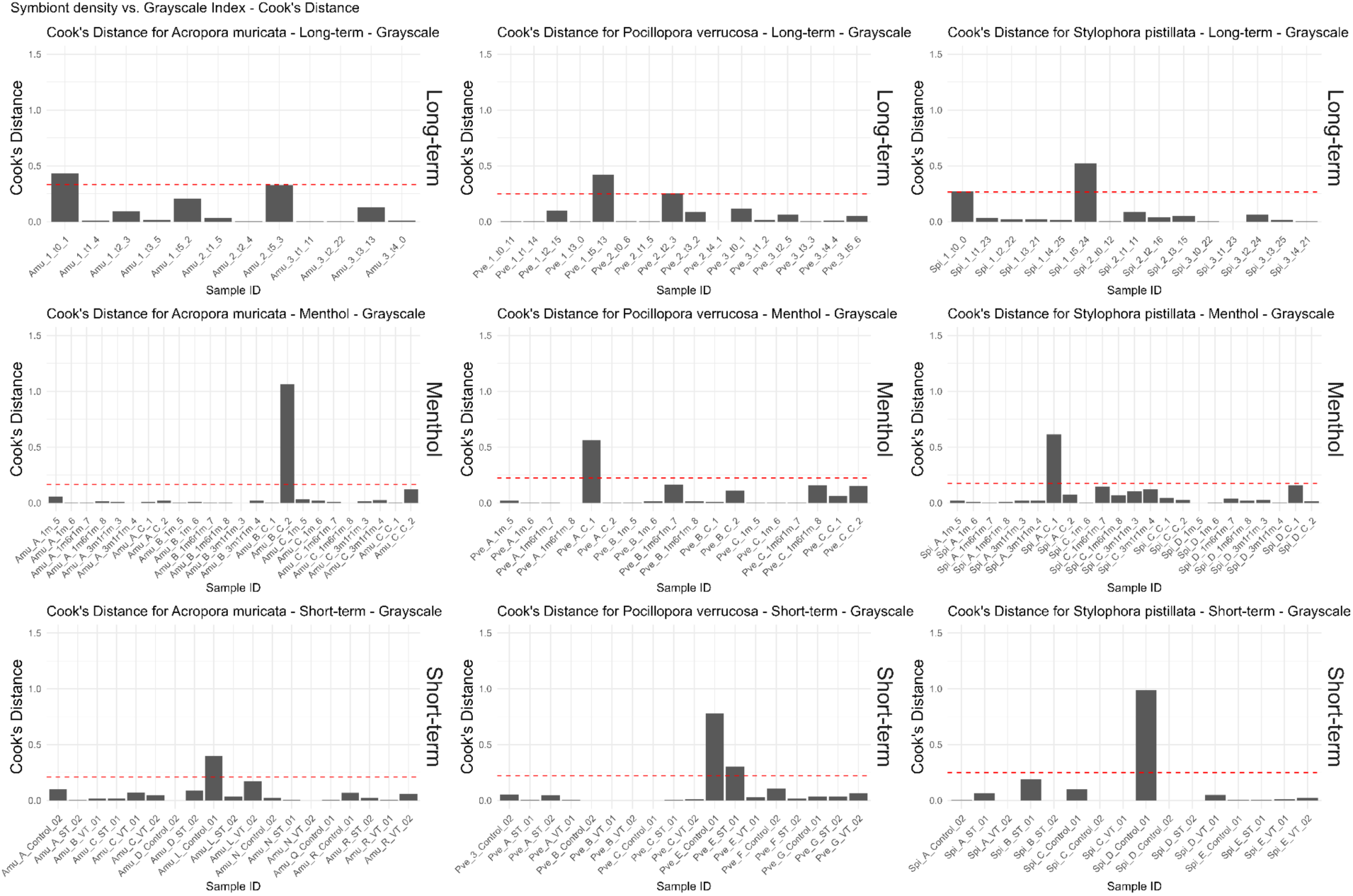
– Samples cook’s distance. **(D**_i_) in the regression model between symbiont cell density and grayscale index across bleaching protocols and coral species. D_i_ is a measure used to identify influential observations (outliers) in the dataset that may disproportionately affect the regression results. Only values with a D_i_ > 0.7 were removed.

